# Single Amplified Genome Catalog Reveals the Dynamics of Mobilome and Resistome in the Human Microbiome

**DOI:** 10.1101/2023.12.06.570492

**Authors:** Tetsuro Kawano-Sugaya, Koji Arikawa, Tatsuya Saeki, Taruho Endoh, Kazuma Kamata, Ayumi Matsuhashi, Masahito Hosokawa

**Affiliations:** bitBiome, Inc., 513 Wasedatsurumaki-cho, Shinjuku-ku, Tokyo, 162-0041, Japan; Department of Life Science and Medical Bioscience, Waseda University, 2-2 Wakamatsu-cho, Shinjuku-ku, Tokyo, 162-8480, Japan; Computational Bio Big-Data Open Innovation Laboratory, National Institute of Advanced Industrial Science and Technology, 3-4-1 Okubo, Shinjuku-ku, Tokyo, 169-8555, Japan; Institute for Advanced Research of Biosystem Dynamics, Waseda Research Institute for Science and Engineering, 3-4-1 Okubo, Shinjuku-ku, Tokyo, 169-8555, Japan; Research Organization for Nano and Life Innovation, Waseda University, 513 Wasedatsurumaki-cho, Shinjuku-ku, Tokyo, 162-0041, Japan

**Author notes:** Tetsuro Kawano-Sugaya and Koji Arikawa contributed equally to this work.

## Abstract

The increase in metagenome-assembled genomes (MAGs) has significantly advanced our understanding of the functional characterization and taxonomic assignment within the human microbiome. However, MAGs, as population consensus genomes, often mask heterogeneity among species and strains, thereby obfuscating the precise relationships between microbial hosts and mobile genetic elements (MGEs). In contrast, single amplified genomes (SAGs) derived via single-cell genome sequencing can capture individual genomic content, including MGEs. We present the bbsag20 dataset, which encompasses 17,202 human-associated prokaryotic SAGs and 869 MAGs, spanning 647 gut and 312 oral bacterial species. The SAGs revealed diverse bacterial lineages and MGEs with a broad host range that were absent in the MAGs and traced the translocation of oral bacteria to the gut. Importantly, our SAGs linked individual mobilomes to resistomes and meticulously charted a dynamic network of antibiotic resistance genes (ARGs) on MGEs, pinpointing potential ARG reservoirs in the microbial community.

## Introduction

The intimate connection between humans and their associated microbiomes has received significant research attention given its crucial ramifications, including its influence on human health, disease progression, and treatment responses^1–5^. The advent of metagenomics has provided unprecedented insights, particularly by unlocking data from uncultured microbes. Genome catalogs such as the Unified Human Gastrointestinal Genome Catalogue^6–11^ stand out in this endeavor, curating comprehensive microbial genomes from microbial communities. Predominantly comprising metagenome-assembled genomes (MAGs) and isolate genomes, these catalogs are not without limitations.

Notably, metagenomics, in its principle of assembling and aggregating similar sequences, struggles to render MAGs that link information on highly conserved sequences, such as rRNA genes, and mobile genetic elements (MGEs), including plasmids and phages. The limitations of metagenomics have been previously reported^12–15^. For instance, only 7% of even highly complete human gut MAGs yielded 16S rRNA genes^16^. Furthermore, another study reported low presence rates of MGEs in MAGs (38-44% for genomic islands and 1-29% for plasmids) and a complete lack of virulence genes and antibiotic resistance genes (ARGs) in plasmids^17^.

Single-cell genome sequencing has emerged as a potential avenue to overcome these challenges by constructing single amplified genomes (SAGs) from individual microbial strains, including highly conserved genes and MGEs. While this method theoretically reveals cell-to-cell variation, its practical realization depends on the evolution of supporting technologies. Despite advancements in high-throughput single-cell genome sequencing technologies, such as droplet barcoding sequencing^18–20^ and their ability to concurrently acquire tens of thousands of SAGs, several challenges persist. These include low completeness of SAGs, often within the range of 0.01-10%, and the subsequent necessity of pooling numerous SAGs or integration with metagenomes to construct consensus genomes. This limitation results in the recovery of fewer than 100 representative genomes from tens of thousands of sparse SAGs and carries the risk of obscuring strain heterogeneity information, such as the relationship between the host and MGE or ARG.

We have previously developed a single-cell genome sequencing technology with quality and throughput, named SAG-gel^21,22^, which enables the simultaneous generation of hundreds or thousands of SAGs. It can obtain SAGs above medium quality without pooling the SAGs to generate consensus genomes. This advantage is attributed to efficient whole-genome amplification and deep single-cell sequencing by coupling in-gel and well-formatted reactions. Thus far, we have applied our method to various microbiomes, not only from human-associated samples but also from environmental samples, enabling us to reach novel implications such as strain heterogeneity, including MGEs^14,21–24^.

In this study, we aimed to delve deeply into human-associated microbiomes using single-cell genome sequencing and demonstrate the advantages of SAGs in understanding the functions of commensal bacteria. To this end, we present the bbsag20 dataset, which comprises 17,202 SAGs of medium quality and above derived from the human oral and gut microbiomes of Japanese individuals using SAG-gel technology. This dataset, which is one of the largest human oral and gut bacterial SAGs, offers a rich resource for exploring the intricate dynamics of the microbiomes, mobilomes, and resistomes. Through meticulous evaluation of the quality and taxonomic consistency across metagenomes, MAGs, and SAGs, we uncovered compelling evidence of oral bacterial translocation to the gut at the cellular level. Furthermore, our exploration of the mobilome and resistome of gut bacteria, based on cell-resolved SAGs, elucidated unexpectedly broad host ranges of plasmids and phages and detailed individual differences in ARG and MGE prevalence and their networks that are difficult to discern by metagenomics alone.

## Results

### Comparison of genomes obtained by metagenomics and single-cell genomics

The workflow and an overview of the bbsag20 dataset are shown in Fig. 1. Briefly, we performed single-cell genome sequencing^21^ of saliva and fecal samples collected from the Japanese participants (Supplementary Data Table 1). From 32 saliva samples, we obtained 11,809 bacterial SAGs, with an average of 369 SAGs (66 species) per sample. From 51 fecal samples, we obtained 19,042 bacterial SAGs, with an average of 373 SAGs (54 species) per sample. For the same set of 51 fecal samples, shotgun metagenome sequencing was performed, which yielded 1,544 fecal MAGs with an average of 30 MAGs (26 species) per sample (Fig. 1a). Of the 30,851 SAGs, 17,202 (55.76%) were classified as high- or medium-quality. In contrast, 869 of 1,544 MAGs (56.28%) met this criterion (Fig. 1a, b and Supplementary Data Table 2). When examining shared sequence information, SAG contigs shared, on average, a 49.5% overlap with metagenome assembly contigs, ranging between 31.9% and 84.0% (Supplementary Data Fig. 1). Conversely, the overlap with MAGs averaged 30.6%, ranging from 6.9% to 80.4%. Although the commonality of sequences obtained by metagenomics and single-cell genomics depends on the sample, more than half of the sequences were obtained in a method-dependent manner. These disparities underscore the unique genomic information yielded by single-cell genomics compared with metagenomics.

**Fig. 1.**
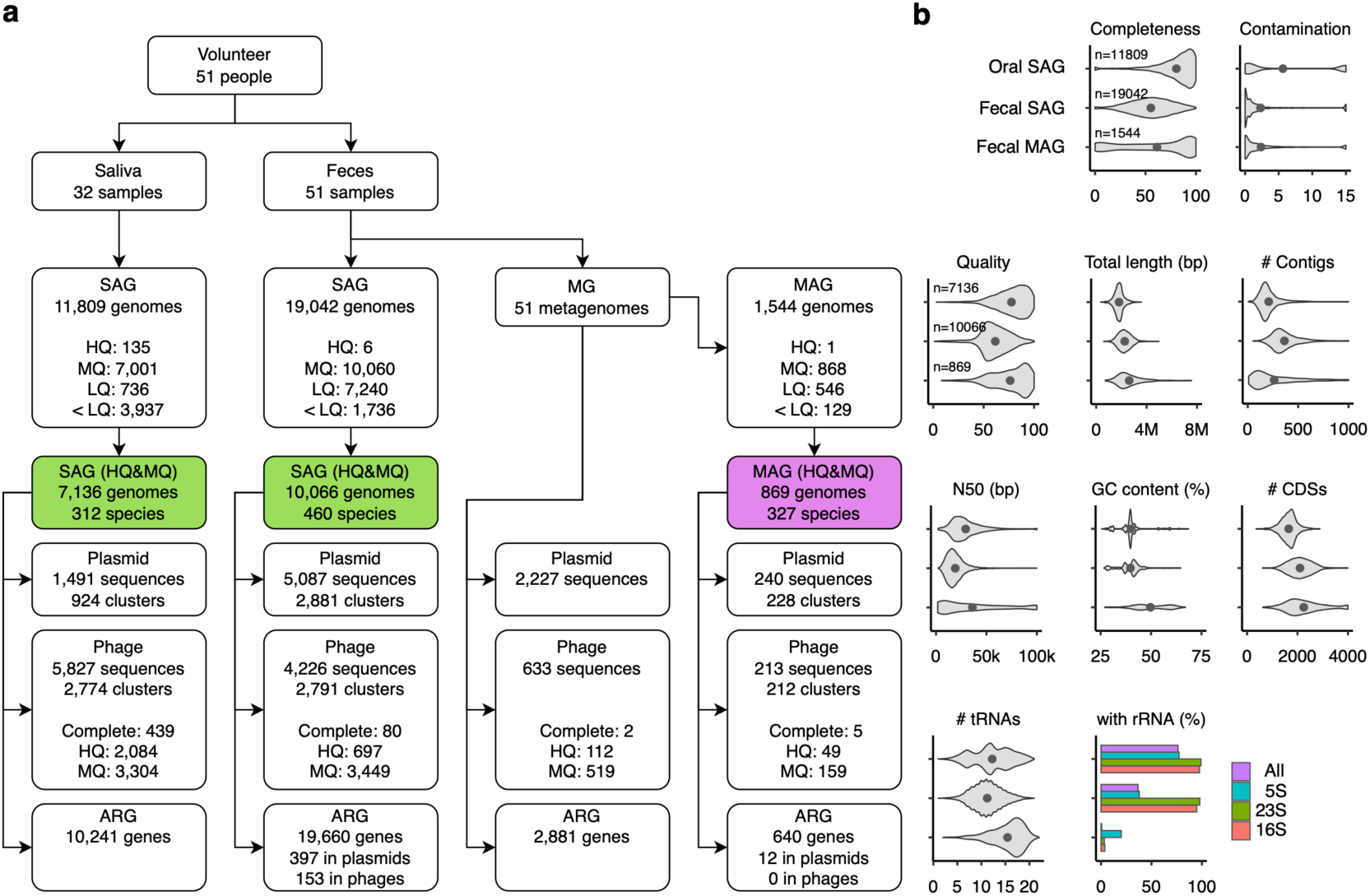
Overview of the Single Amplified Genome catalog bbsag20 for human oral and fecal bacteria. **a**, Overview of samples, assembled genomes, MGEs, and ARGs in the bbsag20 dataset. SAGs and MAGs were categorized as high-quality (HQ), medium-quality (MQ), or low-quality (LQ). **b**, Assembly statistics for both SAGs and MAGs. Gray dots indicate the average values. Genome completeness and contamination show all oral SAG, fecal SAG, and MAG data. Metrics for high- or medium-quality genomes include quality (defined as completeness minus 5x contamination), total length, contig count, N50, GC content, CDSs, tRNAs, and rRNAs.

Comparisons of genome quality (Fig. 1b) showed that high- or medium-quality SAGs tended to have slightly lower quality (mean 61.5), higher contig counts (mean 364.2), and fewer tRNA genes than MAGs. A striking difference was observed in the recovery of rRNA genes, with MAG containing almost no rRNA (0.0069%), whereas 94.8% of fecal SAGs contained 16S rRNA genes, and 36.6% contained full-set rRNA genes (Fig. 1b). This lack of rRNA sequence challenge resulted in the production of a large number of semi-HQ MAGs, representing over a quarter of all MAGs, marked by the absence of rRNA genes, yet showing >90% completeness and <5% contamination. Importantly, while SAG datasets may harbor genomic duplicates of identical strains, MAGs tend to reflect consensus genomes from individual metagenomic samples. Participant-wise species distributions revealed 25–77 (mean 45) species in oral SAGs, 3–54 (mean 30) species in fecal SAGs, and 2–38 species (mean 17) in fecal MAGs. Notably, a Crohn’s disease patient had almost all SAGs (326 of 328) attributed to *Clostridium perfringens*, which causes gas gangrene and enterotoxemia.

A phylogenetic tree, reconstructed from the taxonomy along 10,066 fecal SAGs and 869 MAGs using phyloT (https://phylot.biobyte.de), displayed varying taxonomic biases between SAGs and MAGs (Fig. 2a). A majority of SAGs identified deep genomic diversity across related species in specific lineages, whereas MAGs covered a broader range of lineages. In particular, 96.4% of SAGs targeted 419 species of Firmicutes, currently renamed Bacillota, which are largely absent in MAGs (Fig. 2a; green strips). Of the 460 SAG species identified, 320 were exclusive to SAGs, constituting 49.5% of the combined 647 species from fecal SAGs and MAGs (Fig. 2b and Supplementary Data Table 2). In contrast, MAGs identified 327 species (Fig. 2a; magenta strips), some of which were uncharted in the SAG datasets. (Fig. 2b; 187 species). The predominance of Firmicutes (Gram-positive) in fecal SAGs was similar to that observed in our previous study^14^. These observations could result from inherent sample biases or potentially because certain species, such as gram-negative bacteria, are susceptible to aerobic sample processing, solvent-induced lysis during sample preservation^24,25^, and freezing-induced stress, impeding their recovery through single-cell genome sequencing. Given that single-cell genomics can rectify the phylogenetic biases overlooked in metagenomics and provide strain genomes of closely related species, jointly leveraging both techniques promises a comprehensive genomic reference to unravel microbial diversity.

**Fig. 2.**
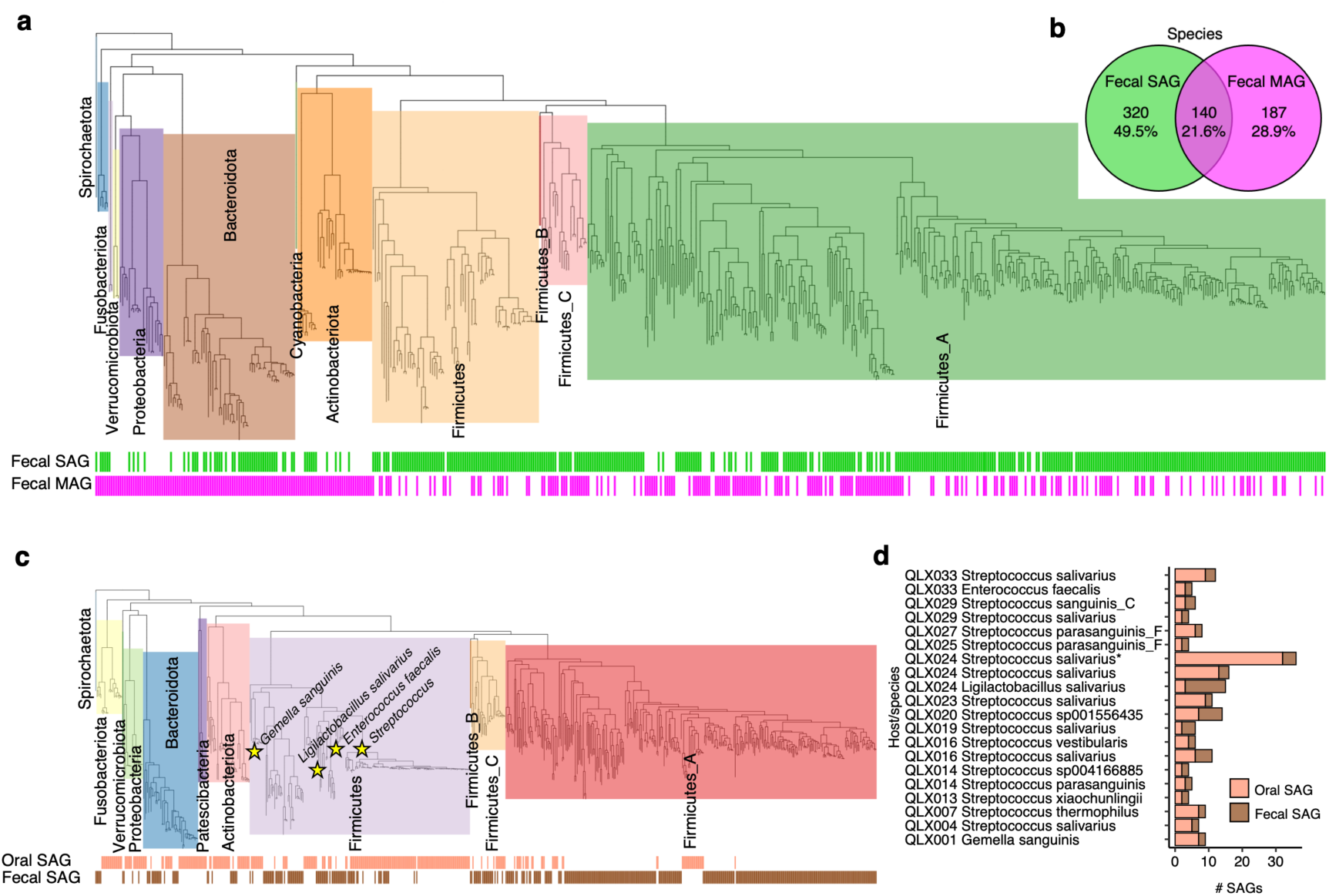
Taxonomy of bbsag20 for human oral and fecal bacteria. **a**, Phylogenetic tree representing the taxonomy of 10,066 fecal SAGs and 869 MAGs. The bottom colored strips show the presence of genomes in each method. **b**, Venn diagram visualizing the species found by fecal SAGs versus MAGs. **c**, Phylogenetic tree detailing the taxonomy of 7,136 oral SAGs compared to 10,066 fecal SAGs. The bottom colored strips show the presence of genomes in each sample. The genera shared between oral and fecal SAGs are marked by stars. **d**, A list of the 12 species consistently present in both the oral and fecal microbiomes of the participants. The number of SAGs obtained is shown by oral and fecal samples in different colors.

### Cell-resolved SAGs revealed oral-to-gut bacterial translocation

The oral microbiome comprises over 700 species and has been implicated in various systemic diseases^26^, including afflictions of the central nervous system, gastrointestinal system, respiratory system, and hypertension^27^. While recent research suggests that 125 out of 310 oral species can be found in both the saliva and feces of 470 individuals across five countries, as determined by shotgun metagenome sequencing^28^, there exists a contrasting study challenging the colonization of oral bacteria in the gut^29^. Techniques, such as shotgun metagenomics and 16S rRNA gene amplicon sequencing, primarily capture genomic fragment information. These methods might overstate the extent of oral bacterial translocation to the gut, especially because they also detect DNA fragments from lysed cells.

To investigate the translocation of oral bacteria to the gut at the cellular level, we analyzed the taxonomy of SAGs from both saliva (7,136 SAGs across 312 species) and feces (10,066 SAGs across 460 species) for each participant (Fig. 2c). The overlap between these two microbiomes was limited, with only 12 species from four genera in oral SAG detected in fecal SAGs (Fig. 2c; stars). These included *Streptococcus* (nine species), *Enterococcus* (one species), *Ligilatobacillus* (one species), and *Gemella* (one species). Fig. 2d provides details such as sample IDs, species names, and counts of the detected SAGs. Bacterial candidates for translocation were identified based on oral and fecal SAG pairs that showed a Jaccard index >0.21 in Dashing2^30^ within the same participants, suggesting that they represent the same bacterial lineages. Notably, the genus *Streptococcus* exhibited varying species detection trends across participants, and some participants even showed the translocation of multiple species. In total, 14 of the 32 participants, including four who were healthy, displayed signs of translocation. Although we recognize the need for more rigorous validation of gut colonization by oral bacteria, necessitating enhanced gut content sampling, unbiased sample processing, and extensive SAG datasets, our data present initial evidence of bacterial translocation from the oral cavity to the gut based on cell-resolved SAG identity. Utilizing cell-resolved SAGs may be instrumental for culture-independent evaluations of bacterial viability and colonization, especially when exploring the interactions between distinct bacterial species across environments.

### Linking mobilome and resistome in the human-associated microbiome

MGEs, such as plasmids and phages, are transferred across bacterial hosts and sometimes act as carriers of ARGs, thereby conferring antimicrobial resistance to bacteria^31,32^. Despite efforts in culturomics^33,34^ and metagenomics, which have accumulated hundreds of thousands of MGEs^35,36^, current genomic analyses have found it challenging to reveal the prevalence of MGEs in individual bacteria. Unlike traditional methods, SAGs can directly determine the host and MGE relationships based on single-cell-resolved information. To integrate the mobilome and resistome information from SAGs, we detected plasmids using Platon^37^, which matched known databases, and identified phages using PhageBoost^38^. The phages were of complete, high, or medium quality and contained viral genes^39^ obtained from both SAGs and MAGs. From the oral SAGs, we identified 1,491 plasmid sequences and 5,827 phage sequences (Fig. 1a and Supplementary Data Tables 3 and 4). In fecal SAGs, we identified 5,087 plasmid and 4,226 phage sequences, respectively. Oral SAGs tend to have fewer plasmids than fecal SAGs, with 0.21 plasmids/genome compared to 0.51 plasmids/genome. In contrast, oral SAGs contained more phages, with 0.82 phages/genome compared of 0.42 phages/genome. In contrast, of the 2,227 plasmids and 633 phages identified in fecal metagenomes, only 10.78% and 33.65%, respectively, were binned into MAGs, highlighting the challenge of associating MGEs with MAGs. Participant-wise plasmid distributions revealed 2–521 in oral SAG, 4–1331 in fecal SAG, 0–34 in fecal MAG, and 4–130 in fecal MG (Supplementary Data Fig. 2). Participant-wise phage distributions revealed 32–471 in oral SAG, 11–191 in fecal SAG, 0–16 in fecal MAG, and 1–23 in fecal MG samples. The majority (83.1–96.7%) of the oral and fecal phages found were Caudoviricetes, with complete, high-, or medium-quality viral genomes acquired in thousands (Supplementary Data Tables 4).

Next, we evaluated the number of bacterial host lineages and assumed host ranges for each MGE by clustering using MMseqs2^40^ at 90% similarity and coverage. Both plasmids and phages showed distinct broad host ranges when comparing SAGs with MAGs. The histogram showed that 21 species for plasmids and four species for phages were the maximal MGE host ranges observed in fecal SAGs, but only three species for plasmids and one species for phages were observed in fecal MAGs (Fig. 3a). These observations in SAGs are consistent with a recent study of broad-host-range plasmids using thousands of isolated genomes in public databases^41^ and demonstrate the advantage of single-cell genomics for determining the bacterial host ranges of MGEs, which are often underestimated using conventional metagenomic approaches.

**Fig. 3.**
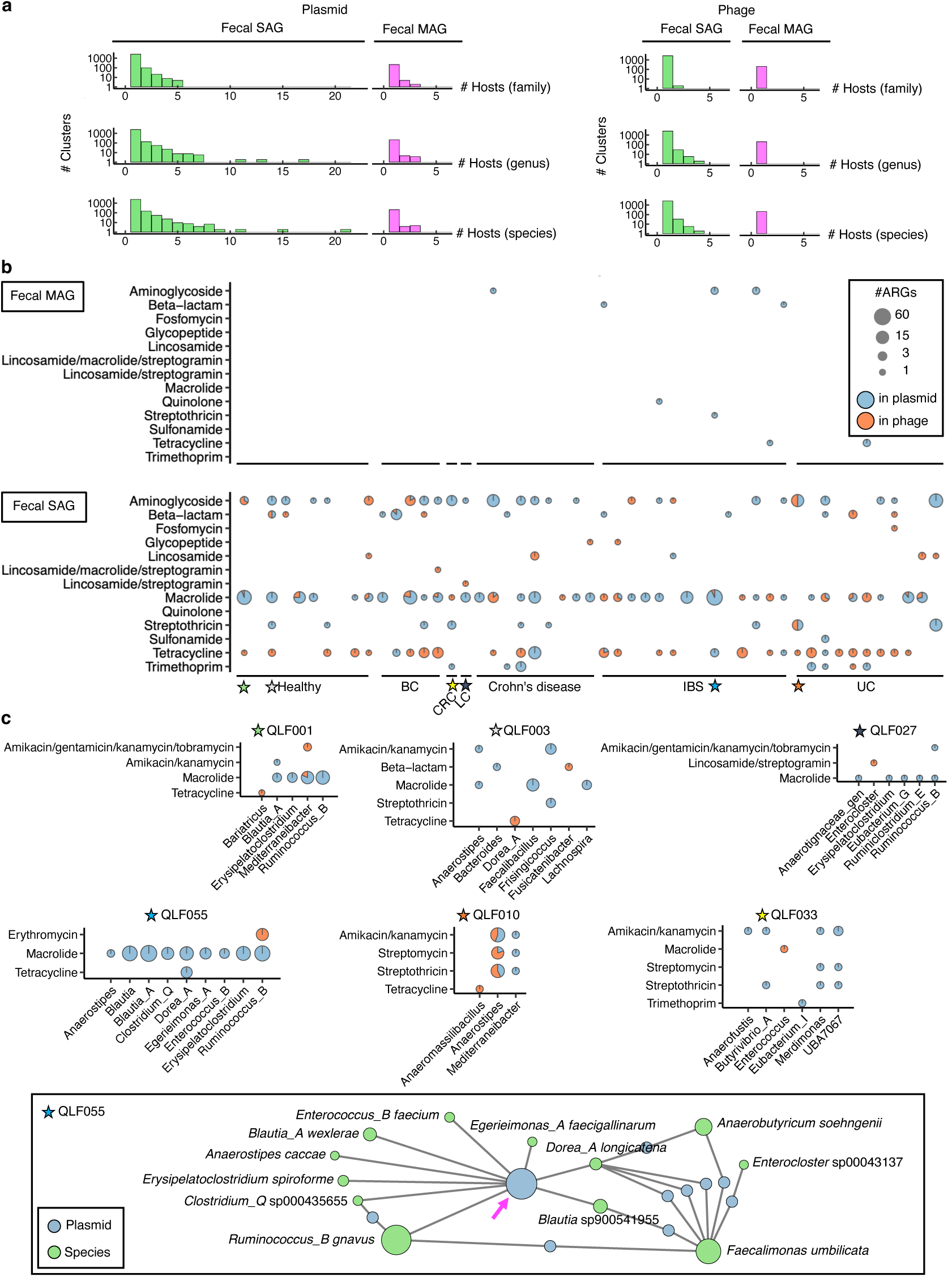
Detailed examination of mobilomes and resistomes in human-associated microbiomes at single-cell resolution. **a**, Determination of the host spectrum of plasmids and phages. To avoid redundant counts, similar plasmids or phage sequences were grouped into clusters. The predicted host numbers are depicted in histograms, distinguishing between SAGs and MAGs across different taxonomic ranks. **b**, Distribution of ARGs in MGEs. ARG (class) presence and genetic context are visualized as pie charts. The x-axis labels detail the medical condition associated with each sample (Healthy; BC, breast cancer; CRC, colorectal cancer; LC, lung cancer; IBS, irritable bowel syndrome; UC, ulcerative colitis). **c**, Comparison of ARGs (subclasses) in MGEs among participants. Six resistomes in the gut microbiome (QLF001, QLF003, QLF010, QLF027, QLF033, and QLF055, marked with stars in **c**) are presented. For QLF055, a network diagram depicted the links between the plasmid and its host genome at the species level. Lines represent the connections between bacterial hosts and plasmids.

We identified 10,241 and 19,660 ARGs in oral and fecal SAGs, respectively (Supplementary Data Table 5), using AMRFinderPlus^42^. Metagenome assemblies displayed 2,881 ARGs, with only 640 allocated to MAGs (Fig. 1a). Notably, fecal SAGs exhibited a higher count of ARGs than MAGs, with 1.95 ARGs/SAG and 0.74 ARGs/MAG. The repertoire of ARGs differed among the oral SAGs, fecal SAGs, metagenome assemblies, and MAGs. The efflux pump genes corresponding to fluoroquinolone resistance were exclusively found in oral SAGs (1,329 of pmrA genes), but not in fecal SAGs or metagenomes (Supplementary Data Fig. 3 and Supplementary Data Table 5; 0 genes and 1 qnrS1 gene, respectively). Conversely, fecal SAGs contained 1,869 genes along with 26 aminoglycoside resistance genes, whereas oral SAGs had only 10 genes (Supplementary Data Fig. 3 and Supplementary Data Table 5). Regarding the disparities between SAGs and MAGs, 631 genes linked to trimethoprim resistance (dfrA1, dfrA17, dfrF, and dfrG) were found in fecal SAGs, while the metagenome and MAG had 21 and 1 genes, respectively. Metagenomes and MAGs showed distinct profiles in tetracycline resistance genes; only 29 genes were found in MAGs, despite 251 genes being found in metagenome assemblies, suggesting difficulty in binning ARGs to MAGs.

Understanding the mode of ARG transfer between bacteria is important for determining the emergence of drug-resistant bacteria. We integrated the mobilome and resistome of fecal SAGs and MAGs to determine the potential for ARG transfer associated with plasmids and phages (Fig. 3b). Importantly, only 2.8% (550/19,660) and 1.8% (12/640) of ARGs in fecal SAGs and MAGs were located on plasmids or phages, respectively, and the rest were found in the chromosome or unidentified (Supplementary Data Fig. 4). There was no obvious dependence of the resistome profiles on the participant background. MAGs detect a minimal number of MGE and ARG relationships, rendering sample comparisons challenging. In contrast, SAGs provided comprehensive data, revealing largely consistent mobilome and resistome profiles across samples (Fig. 3b). This provides an insight into the preferences of the transfer modes for each resistance. For instance, tetracycline resistance genes were mainly found in phages rather than plasmids, whereas those for macrolides were found in plasmids. Intriguingly, although the pattern of resistome possession in each individual was similar, each ARG was shown to be capable of being transferred via plasmids or phages.

Fig. 3c shows the ARG subclasses and their bacterial hosts for the six participants. These bacterial host-specific ARG-MGE profiles suggest that the same resistance genesfor macrolide (ermB) are transmitted via different modes to different bacterial taxa (Fig. 3c; QLF001, 003, 027, and 055). The distribution of resistance genes offers insight into their transmission patterns. For instance, resistance genes for amikacin/kanamycin (aph(3’)-IIIa) predominantly reside on plasmids in QLF033. In QLF010, nearly half of these genes were present in phages of bacterial genera that were not found in QLF033. In other cases, while QLF003 had ermB genes across plasmids from three genera: *Anaerostipes*, *Faecalibacillus*, and *Lachnospira*, QLF055 had these genes across plasmids from nine genera: *Anaerostipes*, *Blautia*, and others (Fig. 3c). The plasmids across the nine genera were identical (90% similarity and coverage), indicating that macrolide resistance genes were transferred via plasmids to various gut bacterial species as reservoirs of ARGs (Fig. 3c; magenta). Single-cell genomics represents a breakthrough in our ability to unveil intricate networks of mobilomes and resistomes on a per-sample basis (Supplementary Data Fig. 5). This information surpasses conventional metagenomics, highlighting dynamic gene exchanges through MGEs in the microbial landscapes of human hosts.

## Discussion

Our study introduced the bbsag20 dataset, which is a comprehensive collection of 17,202 SAGs and 869 MAGs from human saliva and feces. The qualitative similarities between SAGs and MAGs are notable, but the enhanced rRNA gene recovery in SAGs underscores their potential superiority in reference genomes for conventional analyses, including 16S rRNA amplicon sequencing. Both methods exhibited taxonomic biases, emphasizing the benefits of combining single-cell genomics and metagenomics to achieve a full species diversity snapshot. We noted pronounced taxonomic differences between oral and fecal SAGs, with a limited overlap of only 12 species. This mirrors earlier research highlighting the separate microbial niches in the oral cavity and gut, underlining the need for targeted sampling in microbiome studies.

In culture-free microbial research, single-cell genomics has emerged as a potent tool for addressing and filling the lacunae left by traditional metagenomic approaches. This assertion is bolstered by our findings, which highlight the superior sensitivity and precision of single-cell genomics, especially in the profiling of MGEs and ARGs. While SAGs outperformed MAGs in detecting a significantly higher count of MGEs and ARGs, 2.8% of ARGs attributed to plasmids or phages underscored the challenges inherent in identifying ARGs located on MGEs. Single-cell genomics can address the limitations of conventional metagenomics, which struggles to detect ARGs within plasmids and phages. While some cutting-edge research aims to connect MAGs, MGEs, and ARGs using Hi-C metagenomics, the extensive sequence reads required often limit MGE and ARG detection^43–47^. Single-cell genomics, with its ability to overcome such challenges, offers a refined view of the complex dynamics among ARGs, MGEs, and their hosts.

From a public health perspective^48,49^, profiling of the microbiome, mobilome, and resistome highlights pathways to address growing concerns regarding antimicrobial resistance. Recognizing the spread of antimicrobial resistance, it is vital to understand the reservoirs and the transmission of ARGs^3,44,45,50^. This knowledge will drive the development of strategies to prevent the spread of resistant pathogens. For example, discerning that specific resistance genes are mainly present in plasmids within certain bacterial groups may inform both monitoring and targeted interventions.

The proposed research approach has implications not only for health care but also for the environmental and agricultural sectors. With the spread of antimicrobial resistance through diverse ecosystems such as hospitals, farms, and water sources, a thorough understanding of ARG dynamics is essential for a comprehensive approach. Single-cell genomics has the potential to be a key tool for tracking genetic shifts across environments, enabling proactive measures and data-driven decision-making.

In summary, our study emphasizes the game-changing capacity of single-cell genomics in microbiome studies. This provides a new perspective on microbial communities, MGEs, and antimicrobial resistance patterns, and offers a renewed understanding of microbial interplay. The bbsag20 dataset demonstrates the effectiveness of this method. Our data highlight the potential of single-cell genomics for monitoring the dynamics of mobile genetic elements and antibiotic resistance genes in the microbiome across people, animals, and the environment.

## Methods

### Experimental design and sample collection

All human subjects signed a written informed consent form, and the project was approved by the ethics review committee at Yamauchi Clinic (No. 2020-08-00092). All methods were conducted in accordance with the guidelines and regulations outlined by the ethics approval. Preserved feces were collected in 15 mL vials containing 3 mL GuSCN solution (FS-0002; TechnoSuruga Laboratory Co., Ltd., Shizuoka, Japan) and stored at 4°C for a maximum of 2 weeks prior to single-cell encapsulation in droplets or DNA extraction. Preserved saliva was collected in OMNIgene ORAL (OM-501; KYODO INTERNATIONAL INC., Kanagawa, Japan) and stored at 4°C for a maximum of two weeks prior to single-cell encapsulation in droplets or DNA extraction.

### Single-cell genome sequencing

Following the suspension of human feces in the GuSCN solution (500 μL), the supernatant was recovered by centrifugation at 2,000×*g* for 30 s, followed by filtration through a 35-μm nylon mesh and centrifugation at 8,000×*g* for 5 min. The resulting cell pellets were suspended in DPBS and centrifuged twice at 8,000×*g* for 5 minutes. Bacterial cell suspensions were prepared in 100–500 μL of PBS and used in the following steps.

Single-cell genome amplification was performed using the SAG-gel platform, as described in our previous reports. Prior to single-cell encapsulation, cell suspensions were adjusted to 0.3–0.4 cells/droplets in 1.5% agarose in DPBS to prevent encapsulation of multiple cells in single droplets. Using an On-chip Droplet Generator (On-chip Biotechnologies Co., Ltd., Tokyo, Japan), single bacterial cells were encapsulated in droplets and collected in a 1.5 mL tube, which was chilled on ice for 15 minutes to form the gel matrix. Following solidification, the collected droplets were broken using 1H, 1H, 2H, 2H-perfluoro-1-octanol (Sigma-Aldrich, STL, MO, USA) to collect the capsules. The gel capsules were washed with 500 μL of acetone (FUJIFILM Wako Pure Chemical Corporation, Osaka, Japan), and the solution was mixed vigorously and centrifuged. The acetone supernatant was removed, 500 μL of isopropanol (FUJIFILM Wako Pure Chemical Corporation) was added, and the solution was mixed vigorously and centrifuged. The isopropanol supernatant was removed, and the gel capsules were washed three times with 500 μL of DPBS. Individual cells in capsules were then lysed by submerging the gel capsules in lysis solutions: first, 50 U/μL Ready-Lyse Lysozyme Solution (Lucigen, WI, USA), 2 U/mL Zymolyase (Zymo Research Corporation, CA, USA), 22 U/mL lysostaphin (Sigma-Aldrich), and 250 U/mL mutanolysin (Sigma-Aldrich) in DPBS at 37°C overnight; second, 0.5 mg/mL achromopeptidase (FUJIFILM Wako Pure Chemical Corporation) in PBS at 37°C for 6– 8 hours; and third, 1 mg/mL Proteinase K (Promega Corporation, WI, USA) with 0.5% SDS in PBS at 40 °C overnight. At each reagent replacement step, the gel capsules were washed three times with DPBS and subsequently resuspended in the next solution.

Following lysis, the gel capsules were washed five times with DPBS and the supernatant was removed. The capsules were then suspended in Buffer D2 and subjected to multiple displacement amplification (MDA) using REPLI-g Single Cell Kit (QIAGEN, Germany). Following MDA treatment at 30°C for 3 h, the gel capsules were washed three times with 500 μL of DPBS. Thereafter, the capsules were stained with 1 × SYBR Green I (Thermo Fisher Scientific, MA, USA) in DPBS and observed with fluorescence microscopy BZ-X810 (KEYENCE CORPORATION, Osaka, Japan) to count the number of fluorescence-positive gel capsules. Following confirmation of DNA amplification based on the presence of green fluorescence in the gel, fluorescence-positive capsules were sorted into 384-well plates using a BD FACSMelody cell sorter (BD Biosciences, Tokyo, Japan) equipped with a 488-nm excitation laser.

Following droplet sorting, 384-well plates were subjected to the second round of MDA or were stored at −30°C. Following gel capsule collection in 384-well plates, second-round MDA treatment was performed using the REPLI-g Single Cell Kit. Buffer D2 was added to each well and incubated at 65°C for 10 minutes. Thereafter, the MDA mixture was added and incubated at 30°C for 120 minutes. The MDA reaction was terminated by heating at 65°C for 3 minutes.

For sequencing analysis, sequencing SAG libraries were prepared from the second-round MDA product using QIAseq FX DNA Library Kit (QIAGEN). Aliquots of SAGs were transferred to replica plates for DNA yield quantification using Quant-iT dsDNA Broad-Range (BR) Assay Kit (Thermo Fisher Scientific) prior to library preparation. Ligation adaptors were modified using TruSeq-Compatible Full-length Adapters UDI (Integrated DNA Technologies, Inc., IW, USA). Each SAG library was sequenced using an Illumina HiSeq X Ten System with a 2 × 150 bp configuration at Macrogen Japan Corp. (Tokyo, Japan) or using an Illumina NextSeq 2000 System with a 2 × 150 bp configuration.

### Shotgun metagenome sequencing

The QIAamp PowerFecal Pro DNA Kit (QIAGEN) was used for total DNA extraction from the saliva and fecal samples. Metagenomic sequencing libraries were constructed from extracted DNA samples with 10 μL (1/5 volume) reactions using the QIAseq FX DNA Library Kit (QIAGEN). Each metagenomic sequencing library was sequenced using the Illumina NextSeq 2000 System 2 x 150 bp configuration.

### Genome analysis

Adapter sequences and low-quality reads were eliminated from raw sequence reads of metagenome sequences and single-cell genome sequences using bbduk.sh (version 38.90; https://sourceforge.net/projects/bbmap/) with following options (qtrim=r trimq=10 minlength=40 maxns=1 minavgquality=15). These quality-controlled reads of single-cell genomes were assembled de novo into contigs using SPAdes (v3.14.0)^51^ with the following options (--sc --careful --disable-rr --disable-gzip-output). Contigs shorter than 1000 bp were excluded from the SAG assemblies. Metagenome reads were assembled using SPAdes with the following options (--meta). MAGs were constructed using three binning tools, including CONCOCT (v1.0.0)^52^, MaxBin 2(v2.2.6)^53^, and MetaBAT 2 (v2.12.1)^54^, with default options, and DAS_Tool (v1.1.2)^55^ was used to refine the binning results. CDSs, rRNAs, and tRNAs were predicted from the SAGs and MAGs using Prokka (v1.14.6)^56^ with the following options (--rawproduct). The completeness and contamination of SAGs and MAGs were evaluated using CheckM (v1.1.2)^57^ lineage workflow with default options. Taxonomy identification was performed using GTDB-Tk (v2.1.0)^58^ with default options, and GTDB release 207.

### Alignment of metagenome assemblies and single-cell genome assemblies

The contig overlap lengths between metagenome assemblies and SAGs were calculated based on the results of BLASTn with the following options (-outfmt 6 - num_threads 4 -perc_identity 95 -max_target_seqs 50000). Only hits above 1000 bp and 99% similarity were extracted using the awk command (awk ’{if($3 >= 99 && $4>=1000) print $0}’). The redundancy was removed by piling up the overlap hits using awk and BEDTools^59^ (cut -f 2,9,10 input.tsv | sort | uniq | awk ’{if($2>$3) print $1 "\t" $3-1 "\t" $2 "\t." "\t0" "\t+"; else if($2<$3) print $1 "\t" $2-1 "\t" $3 "\t." "\t0" "\t+"; }’ | sort - k1,1V -k2,2V -k3,3V | uniq | bedtools merge -i - | awk ’BEGIN{OFS="\t"}{$4 = $3-$2; print $0}’ | sed "1i contig\tstart\tend\tlength").

### Phylogenetic analysis of fecal bacterial genomes

A total of 10,066 fecal SAGs and 869 fecal MAGs of medium quality were retrieved from the bbsag20 dataset. The undetermined taxa in GTDB-Tk (release 207) were removed, and 561 unique taxa were used in the following analysis. The phylogenetic tree was obtained using phyloT with the removal of one species (*Methanobrevibacter_A smithii*) due to an error in phyloT. Tree visualization and annotation were performed using an R package ‘ggtree’^60^.

### Co-existence of bacterial taxa between oral SAGs and fecal SAGs

A total of 7,136 oral and 10,066 fecal SAGs above medium quality were retrieved from the bbsag20 dataset. The undetermined taxa in GTDB-Tk (release 207) were removed, and 638 unique taxa were used in the following analysis. A phylogenetic tree was constructed using phyloT. Tree visualization and annotation were performed using the R package ‘ggtree’.

### Identification of plasmid, phage, and ARGs

SAGs (oral:7,136; feces:10,066), fecal metagenome assemblies (n=51), and MAGs (n=869) above medium quality were used for mobilome and resistome analysis. Plasmids were predicted using Platon (version 1.6)^37^ with default parameters (platon --db ${platondb} --output ${sampleid} --verbose --threads ${cpus} ${fna}). The list was filtered with “#Plasmid Hits” = 1 (True). Phages were predicted using PhageBoost (version 0.1.7)^38^ with default parameters (PhageBoost -f ${fna} -o ${sampleid} --threads ${cpus}), and their quality were assessed using CheckV (v1.0.1)^39^ with following command (checkv end_to_end -d checkv-db-v1.4 -t 4 ${input}.fna result/checkv/${input}). Only hits with checkv_quality = Medium-quality, High-quality, or Complete having at least one viral gene (viral_genes >= 1) were used in the following analysis. We employed clustering on the plasmids and phages by MMseqs2 (version 13.45111)^40^ using a cut-off threshold both of above 90% coverage and similarity (mmseqs cluster --threads ${cpu} --cov-mode 0 -c 0.90 --min-seq-id 0.90 ${mmseqs_db} ${cluster_db} ${cluster_db_tmp}; mmseqs createtsv -- threads ${cpu} ${mmseqs_db} ${mmseqs_db} ${cluster_db} ${sampleid}_c90s90.tsv). ARGs were identified using the NCBI AMRFinderPlus^42^ with following options (amrfinder --plus -p ${faa} -n ${fna} -g ${gff} --threads ${cpus} - a prokka -o ${sampleid}.tsv --nucleotide_output ${sampleid}_amrfp.fna -- protein_output ${sampleid}_amrfp.faa). To exclude virulence genes, heavy metal resistance genes, and partial genes, we removed hits with Method = PARTIALP, PARTIAL_CONTIG_ENDX, PARTIAL_CONTIG_ENDP, PARTIALX, INTERNAL_STOP) and used only hits with Element type = AMR.

### Estimation of host ranges of plasmids and phages

The filtered plasmids and phages data were combined with metadata of high- or medium-quality SAGs or MAGs containing genome ID, sample ID, and GTDB taxonomy (release 207) based on their contig ID. We counted the unique taxa after grouping them by family, genus, or species for each plasmid or phage cluster using the R program. The frequencies of the unique taxa were plotted.

### Visualization of mobilome and resistome in individual hosts

The identified plasmids and ARGs were combined based on contig ID. ARGs in phage genomic regions were extracted using bedtools (sed 1d ${amrfp}.tsv | awk ‘BEGIN {OFS=”\t”}{print $2 “\t” $3-1 “\t” $4 “\tAMRFinderPlus\t.\t” $5}’ > ${sampleid}_amrfp.bed; sed 1,2d ${sampleid}_phageboost.gff | sed “s/gnl|bB|//g” | sed “s/\(QLF…\)/\1sag/g” | awk ‘BEGIN {OFS=”\t”}{print $1 “\t” $4-1 “\t” $5 “\tPhageBoost\t.\t+”}’ > ${sampleid}_phageboost.bed; bedtools intersect -a ${sampleid}_amrfp.bed -b ${sampleid}_phageboost.bed -f 1.00 -wa). The numbers of ARGs in plasmids or phages for each ARG class were counted by sample ID and plotted using the R package ‘scatterpie.’ The number of ARGs in plasmids or phages for each ARG subclass was counted by genus and plotted depending on the sample ID. The network diagram between plasmids and ARGs was generated using the R package ‘igraph’^61^ and visualized using Gephi^62^.

## Supporting information

Supplementary Data 1 Participants metadata

Supplementary Data 2 All genomes 30851SAGs 1544MAGs

Supplementary Data 3 Plasmid list

Supplementary Data 4 Phage list

Supplementary Data 5 ARG list

## Data availability

The raw data produced in this study were deposited at NCBI under BioProject ID PRJNA1030952. The genome assembly and annotation data produced in this study were deposited at FigShare+ (doi: 10.25452/figshare.plus.24473008).

## Acknowledgements

We would like to acknowledge all the participants for their contribution to this study. The recruitment of participants was supported by QLife Inc. (Tokyo, Japan). We thank Ms. Ayako Sasaki, Ms. Kotoe Date, and Ms. Ai Matsushita (bitBiome, Inc.) for technical assistance on single-cell genome sequencing. The super-computing resource was provided by the Human Genome Center, the Institute of Medical Science, and the University of Tokyo. This work was supported by the Tokyo Metropolitan Small and Medium Enterprise Support Center.

## Author contributions

KA, TS, and MH conceived and managed the study. TS, TE, and MH developed the single-cell genome sequencing platform. TS, TE, and AM conducted the genomic experiments and collected data. TKS, KA, and KK constructed a bioinformatic pipeline for assembling metagenomes and single-cell genomes. TKS analyzed the main data. KA and KK provided essential support for TKS. KA and MH supervised the study. TKS and MH wrote the original manuscript. KA and TS supported the writing. All authors have reviewed and approved the final manuscript.

**Correspondence and requests for materials** should be addressed to MH.

## Ethics declarations

Studies involving human participants were reviewed and approved by the Ethics Review Committee of Yamauchi Clinic IRB (Tokyo, Japan). Written informed consent was obtained from all the participants prior to the study.

## Supplementary Information

**Supplementary Fig. 1 Shared sequence information between SAGs, metagenomes, and MAGs.** (left) Overlap of the contigs of fecal metagenomes (gray) and fecal SAGs (green) based on BLASTn. A mean of 49% of the SAG contigs were aligned to the metagenomic contigs. (right) Overlap of the contigs of fecal MAGs (magenta) and fecal SAGs (green). A mean of 30.6% of the SAG contigs were aligned to the MAG contigs.

**Supplementary Fig. 2 Counts and cluster numbers for MGE and ARG.**

**Supplementary Fig. 3 Heatmap for the presence of antibiotic resistance genes in oral SAGs, fecal SAGs, metagenomes, and MAGs.**

**Supplementary Fig. 4 Comparison of the whole ARG profiles of fecal SAG and MAG.**

**Supplementary Fig. 5 Individual mobilome-coding ARGs.**

**Supplementary Data Table 1** Participant information for this study.

**Supplementary Data Table 2** List of SAGs and MAGs derived from the study, including metrics such as genomic cluster ID determined by Dashing2, genome ID, completeness, contamination, quality score, contig count, total genomic length, N50, GC content (%), CDS count, type, sample ID, sample source, and taxonomy based on the GTDB (release 207), counts of tRNAs, rRNAs, plasmids, phages, and ARGs. Species summary for oral SAGs, fecal SAGs, and MAGs are also included.

**Supplementary Data Table 3** List of plasmids identified in metagenomes, MAGs, and SAGs using Platon.

**Supplementary Data Table 4** List of phages detected in metagenomes, MAGs, and SAGs using PhageBoost, further refined by CheckV and classified by geNomad.

**Supplementary Data Table 5** List of ARGs detected in metagenomes, MAGs, and SAGs using AMRFinderPlus.

## References

1. Gilbert, J. A. et al. Current understanding of the human microbiome. Nat. Med. 24, 392–400 (2018).

2. Sepich-Poore, G. D. et al. The microbiome and human cancer. Science 371, (2021).

3. Fredriksen, S., de Warle, S., van Baarlen, P., Boekhorst, J. & Wells, J. M. Resistome expansion in disease-associated human gut microbiomes. Microbiome 11, 166 (2023).

4. Lloyd-Price, J. et al. Multi-omics of the gut microbial ecosystem in inflammatory bowel diseases. Nature 569, 655–662 (2019).

5. Wirbel, J. et al. Meta-analysis of fecal metagenomes reveals global microbial signatures that are specific for colorectal cancer. Nat. Med. 25, 679–689 (2019).

6. Almeida, A. et al. A unified catalog of 204,938 reference genomes from the human gut microbiome. Nat. Biotechnol. 39, 105–114 (2021).

7. Pasolli, E. et al. Extensive Unexplored Human Microbiome Diversity Revealed by Over 150,000 Genomes from Metagenomes Spanning Age, Geography, and Lifestyle. Cell 176, 649–662.e20 (2019).

8. Nayfach, S. et al. A genomic catalog of Earth’s microbiomes. Nat. Biotechnol. 39, 499–509 (2020).

9. Coelho, L. P. et al. Towards the biogeography of prokaryotic genes. Nature 601, 252–256 (2022).

10. Saheb Kashaf, S., et al. Integrating cultivation and metagenomics for a multi-kingdom view of skin microbiome diversity and functions. Nat. Microbiol. 7, 169– 179 (2022).

11. Li, W. et al. A catalog of bacterial reference genomes from cultivated human oral bacteria. NPJ Biofilms Microbiomes 9, 45 (2023).

12. Meziti, A. et al. The Reliability of Metagenome-Assembled Genomes (MAGs) in Representing Natural Populations: Insights from Comparing MAGs against Isolate Genomes Derived from the Same Fecal Sample. Appl. Environ. Microbiol. 87, e02593–20 (2021).

13. Mise, K. & Iwasaki, W. Unexpected absence of ribosomal protein genes from metagenome-assembled genomes. ISME Commun. 2, 118 (2022).

14. Arikawa, K. et al. Recovery of strain-resolved genomes from human microbiome through an integration framework of single-cell genomics and metagenomics. Microbiome 9, 202 (2021).

15. Chen, L.-X., Anantharaman, K., Shaiber, A., Eren, A. M. & Banfield, J. F. Accurate and complete genomes from metagenomes. Genome Res. 30, 315–333 (2020).

16. Hiseni, P., Snipen, L., Wilson, R. C., Furu, K. & Rudi, K. Questioning the quality of 16S rRNA gene sequences derived from human gut metagenome-assembled genomes. Front. Microbiol. 12, 822301 (2021).

17. Maguire, F. et al. Metagenome-assembled genome binning methods with short reads disproportionately fail for plasmids and genomic Islands. Microb. Genom. 6, mgen000436 (2020).

18. Zheng, W. et al. High-throughput, single-microbe genomics with strain resolution, applied to a human gut microbiome. Science 376, eabm1483 (2022).

19. Lan, F., Demaree, B., Ahmed, N. & Abate, A. R. Single-cell genome sequencing at ultra-high-throughput with microfluidic droplet barcoding. Nat. Biotechnol. 35, 640– 646 (2017).

20. Li, X. et al. Microbiome single cell atlases generated with a commercial instrument. bioRxiv (2023) doi:10.1101/2023.08.08.551713.

21. Chijiiwa, R. et al. Single-cell genomics of uncultured bacteria reveals dietary fiber responders in the mouse gut microbiota. Microbiome 8, 5 (2020).

22. Nishikawa, Y. et al. Validation of the application of gel beads-based single-cell genome sequencing platform to soil and seawater. ISME Commun. 2, 1–11 (2022).

23. Ide, K. et al. Exploring strain diversity of dominant human skin bacterial species using single-cell genome sequencing. Front. Microbiol. 13, 955404 (2022).

24. Hosokawa, M. et al. Strain-level profiling of viable microbial community by selective single-cell genome sequencing. Sci. Rep. 12, 4443 (2022).

25. Hosomi, K. et al. Method for preparing DNA from feces in guanidine thiocyanate solution affects 16S rRNA-based profiling of human microbiota diversity. Sci. Rep. 7, 4339 (2017).

26. Deo, P. N. & Deshmukh, R. Oral microbiome: Unveiling the fundamentals. J. Oral Maxillofac. Pathol. 23, 122–128 (2019).

27. Herremans, K. M. et al. The oral microbiome, pancreatic cancer and human diversity in the age of precision medicine. Microbiome 10, 93 (2022).

28. Schmidt, T. S. et al. Extensive transmission of microbes along the gastrointestinal tract. Elife 8, (2019).

29. Rashidi, A., Ebadi, M., Weisdorf, D. J., Costalonga, M. & Staley, C. No evidence for colonization of oral bacteria in the distal gut in healthy adults. Proc. Natl. Acad. Sci. U. S. A. 118, (2021).

30. Baker, D. N. & Langmead, B. Genomic sketching with multiplicities and locality-sensitive hashing using Dashing 2. Genome Res. 33, 1218–1227 (2023).

31. McInnes, R. S., McCallum, G. E., Lamberte, L. E. & van Schaik, W. Horizontal transfer of antibiotic resistance genes in the human gut microbiome. Curr. Opin. Microbiol. 53, 35–43 (2020).

32. Brito, I. L. Examining horizontal gene transfer in microbial communities. Nat. Rev. Microbiol. 19, 442–453 (2021).

33. Lagier, J.-C. et al. Culturing the human microbiota and culturomics. Nat. Rev. Microbiol. 16, 540–550 (2018).

34. Greub, G. Culturomics: a new approach to study the human microbiome. Clin. Microbiol. Infect. 18, 1157–1159 (2012).

35. Lai, S. et al. mMGE: a database for human metagenomic extrachromosomal mobile genetic elements. Nucleic Acids Res. 49, D783–D791 (2021).

36. Schmartz, G. P. et al. PLSDB: advancing a comprehensive database of bacterial plasmids. Nucleic Acids Res. 50, D273–D278 (2022).

37. Schwengers, O. et al. Platon: identification and characterization of bacterial plasmid contigs in short-read draft assemblies exploiting protein sequence-based replicon distribution scores. Microb. Genom. 6, mgen000398 (2020).

38. Sirén, K., et al. Rapid discovery of novel prophages using biological feature engineering and machine learning. NAR Genom. Bioinform. 3, lqaa109 (2021).

39. Nayfach, S. et al. CheckV assesses the quality and completeness of metagenome-assembled viral genomes. Nat. Biotechnol. 39, 578–585 (2021).

40. Steinegger, M. & Söding, J. MMseqs2 enables sensitive protein sequence searching for the analysis of massive data sets. Nat. Biotechnol. 35, 1026–1028 (2017).

41. Yang, L. et al. Global transmission of broad-host-range plasmids derived from the human gut microbiome. Nucleic Acids Res. 51, 8005–8019 (2023).

42. Feldgarden, M. et al. AMRFinderPlus and the Reference Gene Catalog facilitate examination of the genomic links among antimicrobial resistance, stress response, and virulence. Sci. Rep. 11, 12728 (2021).

43. Marbouty, M., Baudry, L., Cournac, A. & Koszul, R. Scaffolding bacterial genomes and probing host-virus interactions in gut microbiome by proximity ligation (chromosome capture) assay. Sci. Adv. 3, e1602105 (2017).

44. Stalder, T., Press, M. O., Sullivan, S., Liachko, I. & Top, E. M. Linking the resistome and plasmidome to the microbiome. ISME J. 13, 2437–2446 (2019).

45. Kent, A. G., Vill, A. C., Shi, Q., Satlin, M. J. & Brito, I. L. Widespread transfer of mobile antibiotic resistance genes within individual gut microbiomes revealed through bacterial Hi-C. Nat. Commun. 11, 4379 (2020).

46. Marbouty, M., Thierry, A., Millot, G. A. & Koszul, R. MetaHiC phage-bacteria infection network reveals active cycling phages of the healthy human gut. Elife 10, (2021).

47. Du, Y., Fuhrman, J. A. & Sun, F. ViralCC retrieves complete viral genomes and virus-host pairs from metagenomic Hi-C data. Nat. Commun. 14, 502 (2023).

48. Baquero, F., Coque, T. M., Martínez, J.-L., Aracil-Gisbert, S. & Lanza, V. F. Gene transmission in the one health microbiosphere and the channels of antimicrobial resistance. Front. Microbiol. 10, (2019).

49. Djordjevic, S. P. et al. Genomic surveillance for antimicrobial resistance - a One Health perspective. Nat. Rev. Genet. (2023) doi:10.1038/s41576-023-00649-y.

50. Berbers, B. et al. Combining short and long read sequencing to characterize antimicrobial resistance genes on plasmids applied to an unauthorized genetically modified Bacillus. Sci. Rep. 10, 1–13 (2020).

51. Bankevich, A. et al. SPAdes: a new genome assembly algorithm and its applications to single-cell sequencing. J. Comput. Biol. 19, 455–477 (2012).

52. Alneberg, J. et al. Binning metagenomic contigs by coverage and composition. Nat. Methods 11, 1144–1146 (2014).

53. Wu, Y.-W., Simmons, B. A. & Singer, S. W. MaxBin 2.0: an automated binning algorithm to recover genomes from multiple metagenomic datasets. Bioinformatics 32, 605–607 (2016).

54. Kang, D. D. et al. MetaBAT 2: an adaptive binning algorithm for robust and efficient genome reconstruction from metagenome assemblies. PeerJ 7, e7359 (2019).

55. Sieber, C. M. K. et al. Recovery of genomes from metagenomes via a dereplication, aggregation and scoring strategy. Nat. Microbiol. 3, 836–843 (2018).

56. Seemann, T. Prokka: rapid prokaryotic genome annotation. Bioinformatics 30, 2068–2069 (2014).

57. Parks, D. H., Imelfort, M., Skennerton, C. T., Hugenholtz, P. & Tyson, G. W. CheckM: assessing the quality of microbial genomes recovered from isolates, single cells, and metagenomes. Genome Res. 25, 1043–1055 (2015).

58. Chaumeil, P.-A., Mussig, A. J., Hugenholtz, P. & Parks, D. H. GTDB-Tk v2: memory friendly classification with the genome taxonomy database. Bioinformatics 38, 5315–5316 (2022).

59. Quinlan, A. R. & Hall, I. M. BEDTools: a flexible suite of utilities for comparing genomic features. Bioinformatics 26, 841–842 (2010).

60. Yu, G., Smith, D. K., Zhu, H., Guan, Y. & Lam, T. T.-Y. Ggtree: An r package for visualization and annotation of phylogenetic trees with their covariates and other associated data. Methods Ecol. Evol. 8, 28–36 (2017).

61. Csardi, G. & Nepusz, T. The igraph Software Package for Complex Network Research. Int. J. Complex Syst. 1695, 1–9, (2006).

62. Bastian, M., Heymann, S. & Jacomy, M. Gephi: An open source software for exploring and manipulating networks. Proc. Int. AAAI Conf. Weblogs Soc. Media 3, 361–362 (2009).

