## Supplementary Information for "Single Amplified Genome Catalog Reveals the Dynamics of Mobilome and Resistome in the Human Microbiome"

### Supplementary Fig. 1

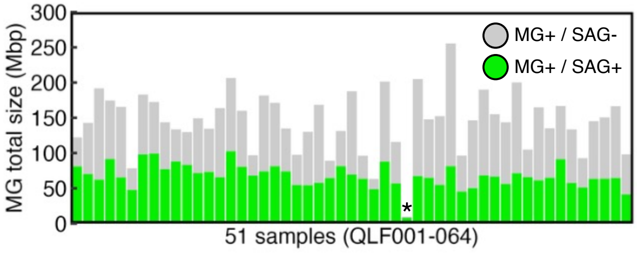


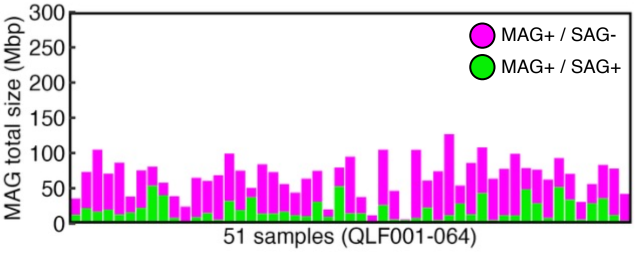

Supplementary Fig. 2

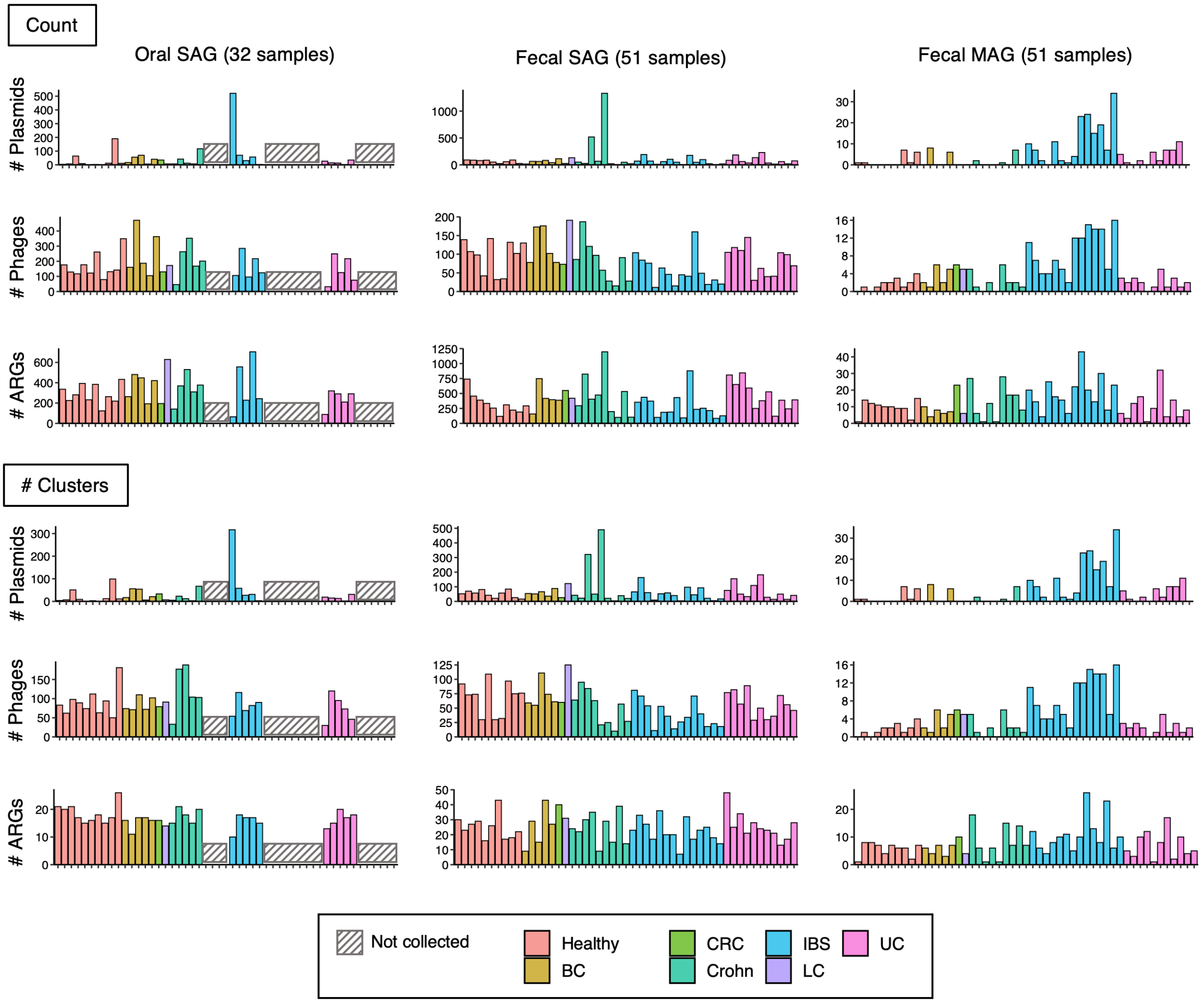

Supplementary Fig. 3

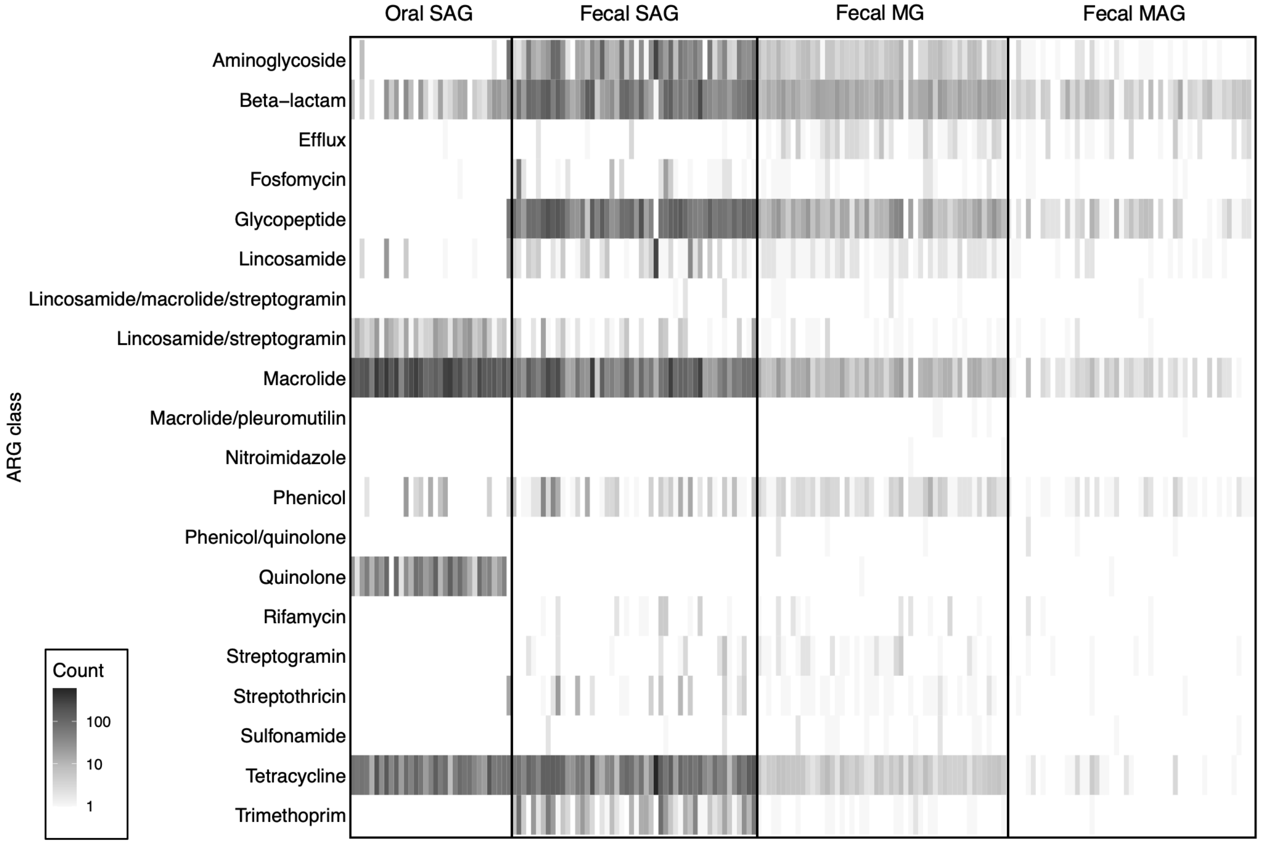

Supplementary Fig. 4

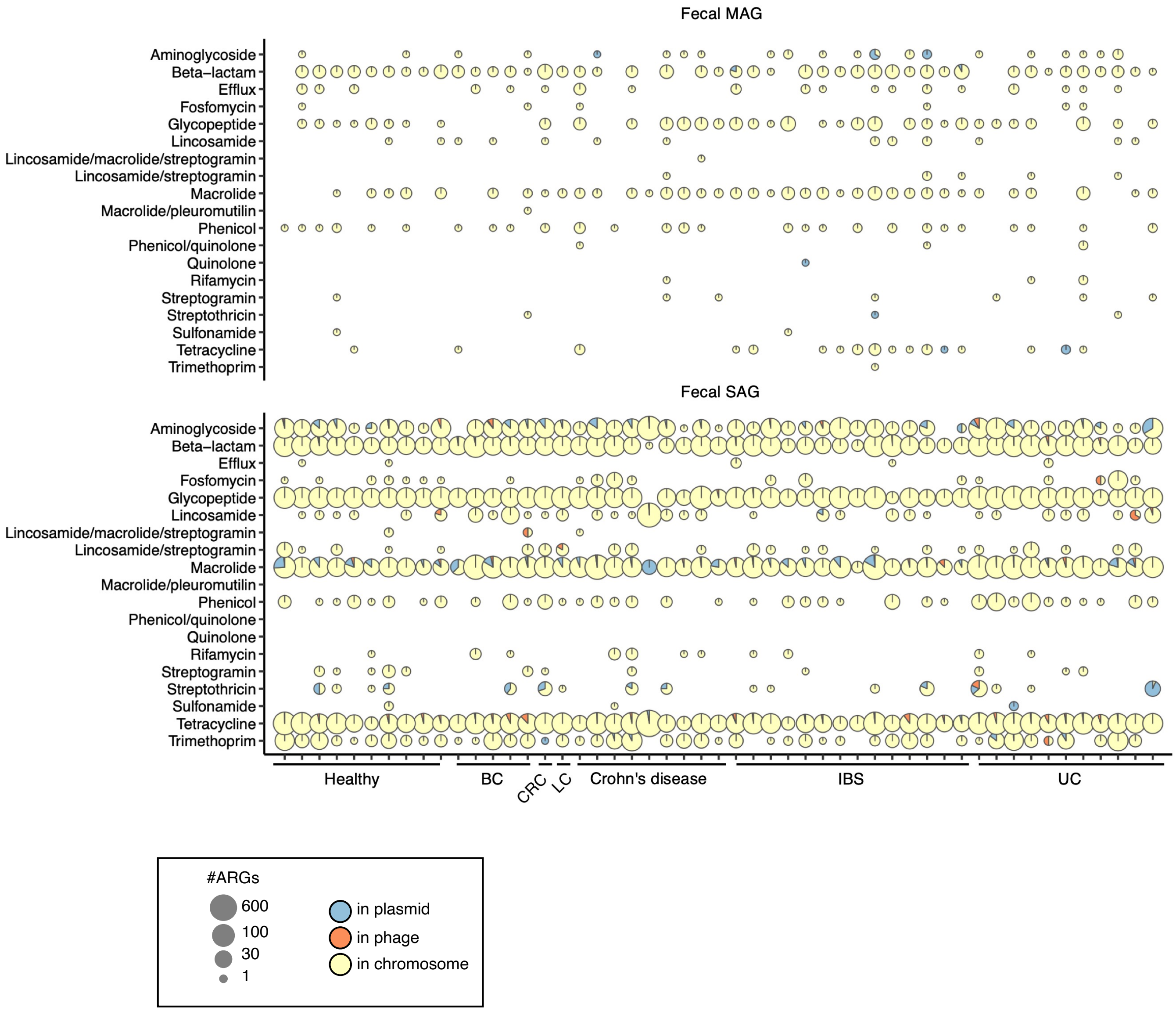

Supplementary Fig. 5 (1)

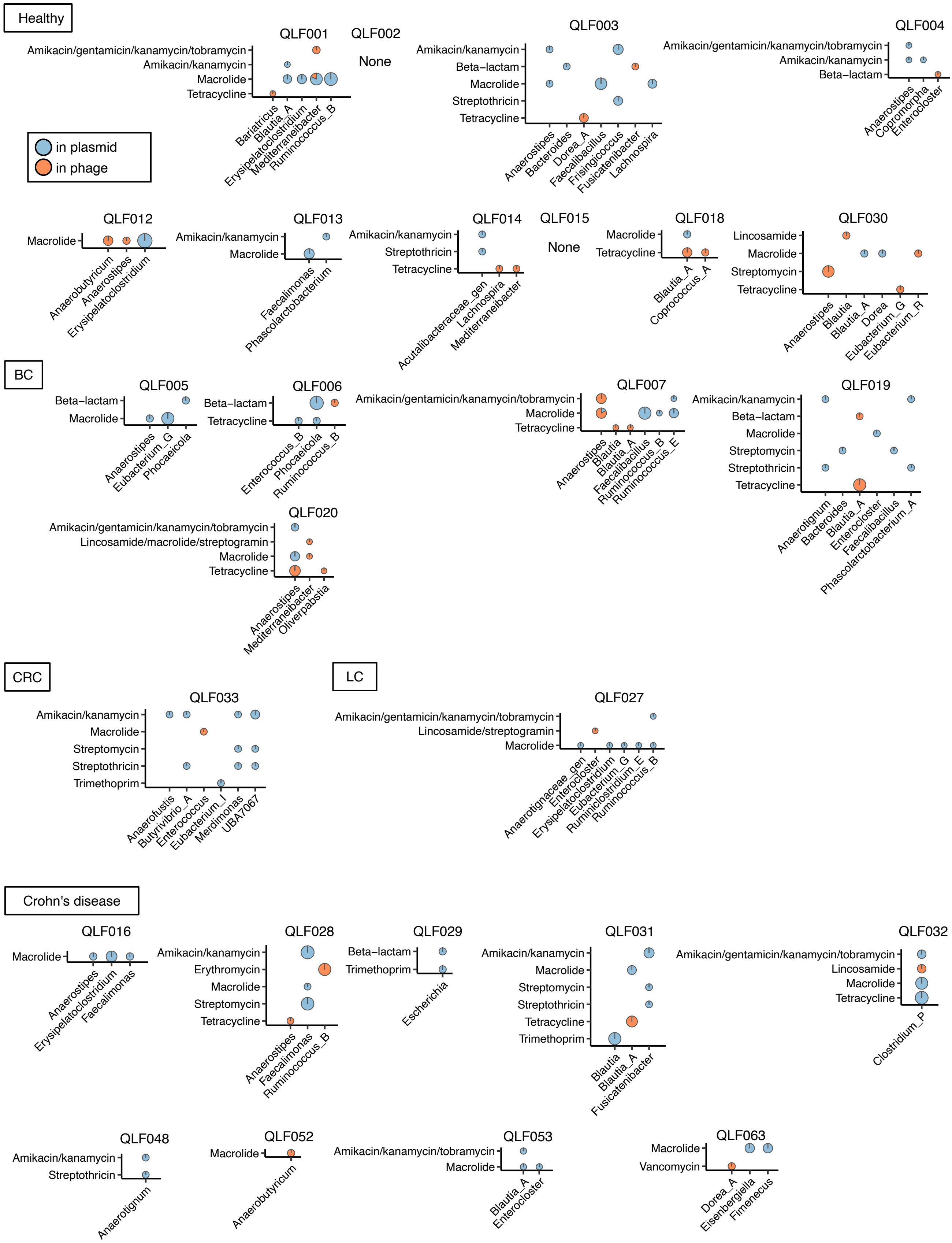

Supplementary Fig. 5 (2)

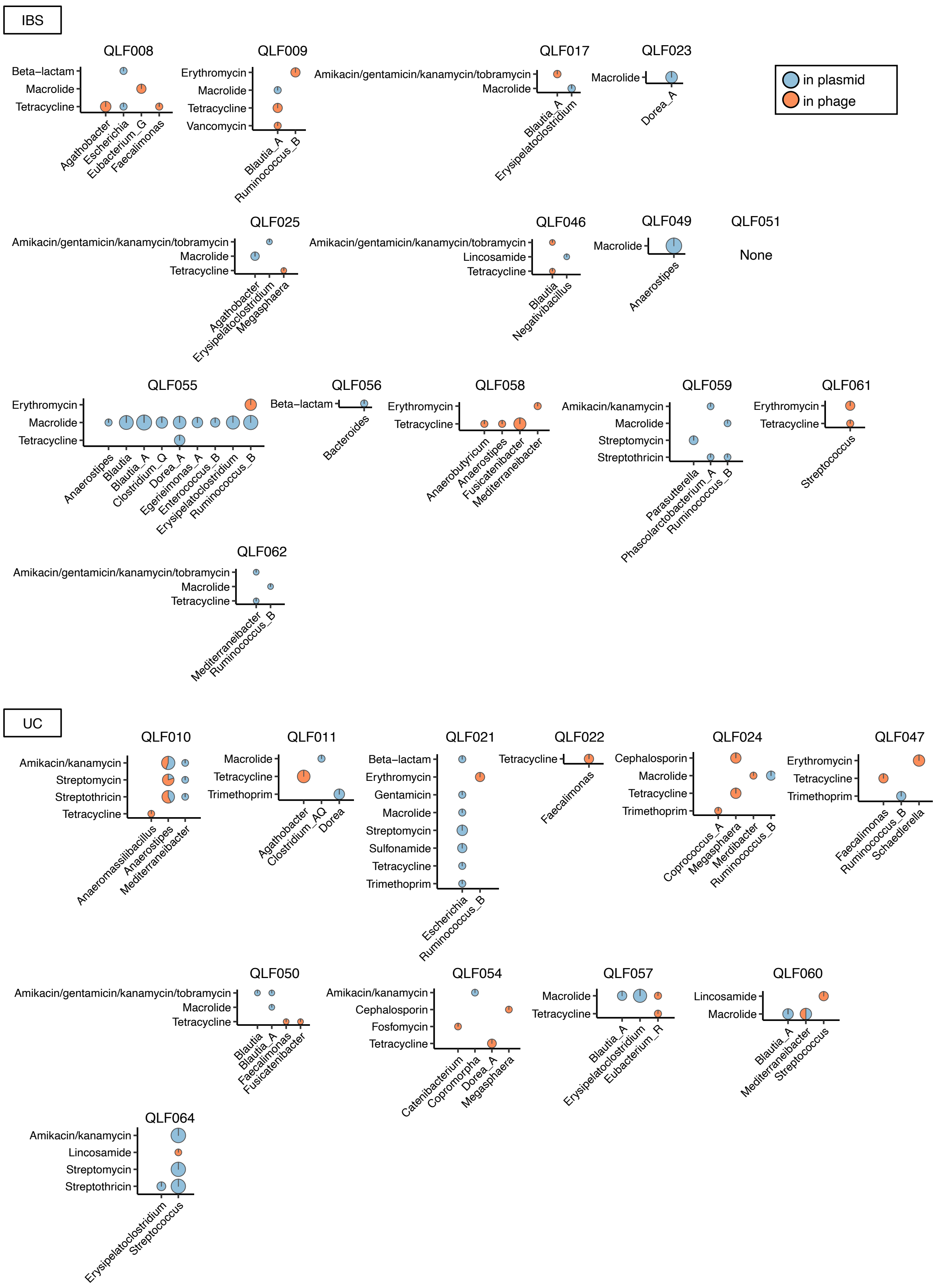
